# Early Embryonic Development of the German Cockroach *Blattella germanica*

**DOI:** 10.1101/2024.07.08.602440

**Authors:** Ariel Bar-Lev Viterbo, Judith R. Wexler, Orel Mayost Lev-Ari, Ariel D. Chipman

## Abstract

**Background:** Early embryogenesis is characterized by dramatic cell proliferation and movement. In most insects, early embryogenesis includes a phase called the uniform blastoderm, during which cells evenly cover the entirety of the egg. However, the embryo of the German cockroach, *Blattella germanica*, like those of many insects within the super order Polyneoptera, does not have a uniform blastoderm; instead, its first cells condense rapidly at the site of a future germband. We investigated early development in this species in order to understand how early gene expression is or is not conserved in these insect embryos with distinct early cell behaviors.

**Results:** We present a detailed time series of nuclear division and distribution from fertilization through germband formation and report patterns of expression for the early patterning genes *hunchback*, *caudal*, and twist in order to understand early polarization and mesoderm formation. We show a detailed time course of the spatial expression of two genes involved in the segmentation cascade, *hedgehog* and *even-skipped,* and demonstrate two distinct dynamics of the segmentation process.

**Conclusions:** Despite dramatic differences in cell distribution between the blastoderms of many Polyneopteran insects and those of more well-studied developmental models, expression patterns of early patterning genes are mostly similar. Genes associated with axis determination in other insects are activated relatively late and are probably not maternally deposited. The two phases of segmentation – simultaneous and sequential – might indicate a broadly conserved mode of morphological differentiation. The developmental time course we present here should be of value for further investigation into the causes of this distinct blastoderm type.

## Introduction

The German cockroach (*Blattella germanica*) has long been a model for post-embryonic development, and its biology has been extensively studied because of its role as a human pest [1]. Nonetheless, there only exists a brief modern description of its embryonic development [2] since Akira Tanaka presented a detailed time series of the animal’s development using light microscopy and cellular dyes in the 1970s [3]. Recent work on the embryonic development of *B. germanica* [2,4] underscores the need to describe the early stages of embryogenesis in this species.

*B. germanica* is a hemimetabolous insect, a member of the superorder Dictyoptera. It has a cosmopolitan distribution and is gregarious. The species is ovoviviparous; *B. germanica* females lay eggs into egg cases (oothecae), which they then carry for the duration of embryonic development (three to four weeks, depending on temperature). The process of depositing eggs into the ootheca takes approximately half a day, leading to a gradient of ages among embryos within a single egg case.

*B. germanica* embryogenesis differs from more well-studied model organisms in a few key ways. First, the organism lacks a uniform blastoderm; that is, there is never a stage in embryogenesis in which nuclei or cells are distributed evenly across the surface of the egg. Instead, the nuclei that will form the germ anlage cluster together in a localized area on the egg’s surface. In 1972, Anderson described blastoderms of this type as direct-differentiating, as opposed to uniform [5]. Uniform blastoderms are found in holometabolan, paleopteran, and hemipteran insect embryos (with exceptions for certain lepidopteran [6,7] and coleopteran [8–10] embryos. Direct-differentiating blastoderms are typical of polyneopteran insect embryos (with exceptions for certain orthopterans [11–13] and plecopterans [14]. Because the preponderance of research into early insect embryogenesis is concentrated in holometabolous model species with uniform blastoderms, there is a relative lack of information about how early patterning proceeds in insects with direct-differentiating blastoderms.

Among those insects with direct-differentiated blastoderms, a subset have embryos that form via the fusion of two regions of high cell density on the lateral sides of the egg [15,16]. The formation of an embryo from such lateral plates is suggested to be an apomorphy of Polyneoptera, a large phylogenetic grouping including roaches, mantises, and crickets [15,16]. The lateral plate fusion that forms the *B. germanica* germband is fundamentally different from the morphogenesis of early development in the other species cited above and has been called “fault type” [17].

In insects, mesoderm formation is tightly linked to specific morphogenetic developments. Mesodermal tissue forms inside of the ventral furrow, which itself is the result of embryonic tissue invaginating. This phenomenon has been extensively reviewed by Anderson [5], Eastham [18], Johannsen and Butt [19], and studied specifically in *Drosophila melanogaster* [20], *Tribolium castaneum* [21] *Gryllus bimaculatus* [22] *Carausius morosus* [23], *Tenebrio molitor* [24] and others. To our knowledge, the expression of the mesodermal marker *twist* in embryos with fault-type mesoderm formation (Polyneoptera) has been investigated only in the cricket *Gryllus bimaculatus* and only in the context of DV patterning [25]. Thus, it is unknown how genetic and morphological cues coordinate to form mesoderm in these insects.

Another notable trait of the *B. germanica* embryo is its lack of blastokinesis, that is the acrobatic movements of an embryo into and then out of the yolk during embryogenesis. While most holometabolous insects and some Hemipterans lack blastokinesis, embryos from all other hemimetabolous insects undergo some sort of movement into and out of the yolk [26]. Even within Blattodea, some taxa have blastokinetic embryos, while others do not. Although embryonic movements and the extra-embryonic membranes that coordinate such movement are beyond the scope of this paper, researchers interested in these questions may find it useful to have a developmental road map for *B. germanica* given the organism’s relatively unusual lack of blastokinesis.

*B. germanica* is a cosmopolitan organism that interacts with humans worldwide, impacting industry and human health. Phylogenetically, it sits in an understudied part of the insect tree, providing potential insights into the evolutionary paths of insect development. Practically, recent work has made CRISPR-Cas9 a viable option for genome editing in the species [27]. For all these reasons, a detailed and modern description of the animal’s embryogenesis is long overdue. The differences between *B. germanica* embryogenesis and that of other, more well-studied insects discussed above mean it is not trivial to simply map events from *B. germanica* embryogenesis onto prior frameworks.

Given *B. germanica*’s phylogenetic position, studying its development can thus shed light on several other evolutionary developmental questions. Prior work from our group has highlighted the correlation between a transition in segmentation mode (simultaneous versus sequential) and final morphology (thorax versus abdomen). As we’ve shown in the well-studied hemimetabolous insect, *Oncopeltus fasciatus,* gnathal and thoracic segments are formed simultaneously [28], while abdominal segments arise sequentially from a segment addition zone [29]. It is unclear whether this link between tagma borders and segmentation modes is specific to *O. fasciatus* (and its relatives) or general to insects. Looking at segmentation in *B. germanica* adds an additional important phylogenetic node to answer this question.

Here, we present a time series of early *B. germanica* embryogenesis, from cleavage stages through segmentation. We show that cell division occurs in pulses in the early blastoderm stages and that cellularization happens at approximately 4-6 percent of development. The early patterning genes *hunchback* and *caudal* are not present in the very early embryo, suggesting they are not maternally deposited. Early stages of mesoderm formation seem similar to those reported in other insects. Segmentation appears to be of the intermediate type – that is, anterior segments are patterned almost simultaneously and posterior ones sequentially – although it is possible to observe the appearance of *hedgehog* stripes one by one.

## Results

### Germband formation

*B. germanica* development takes approximately 25 days under our lab conditions described in the Methods. Germband formation, observed via Sytox and DAPI nuclear staining, takes about 3.5 days, or 14 percent of development (Fig. 1 A-M). We recorded the spatial and temporal dynamics of nuclear divisions during this time. After counting nuclei in embryos collected every 7-8 hours, we noticed two distinct pulses of cell division within the first 65 hours of development (Fig. 1N). These pulses occur between hours 22-30 and 44-53. We noticed that these pulses of cell division corresponded to changes in the spatial distribution of nuclei over the surface of the egg. Combining these temporal and spatial observations, we divided early germband formation into the four following stages.

**Figure 1:**
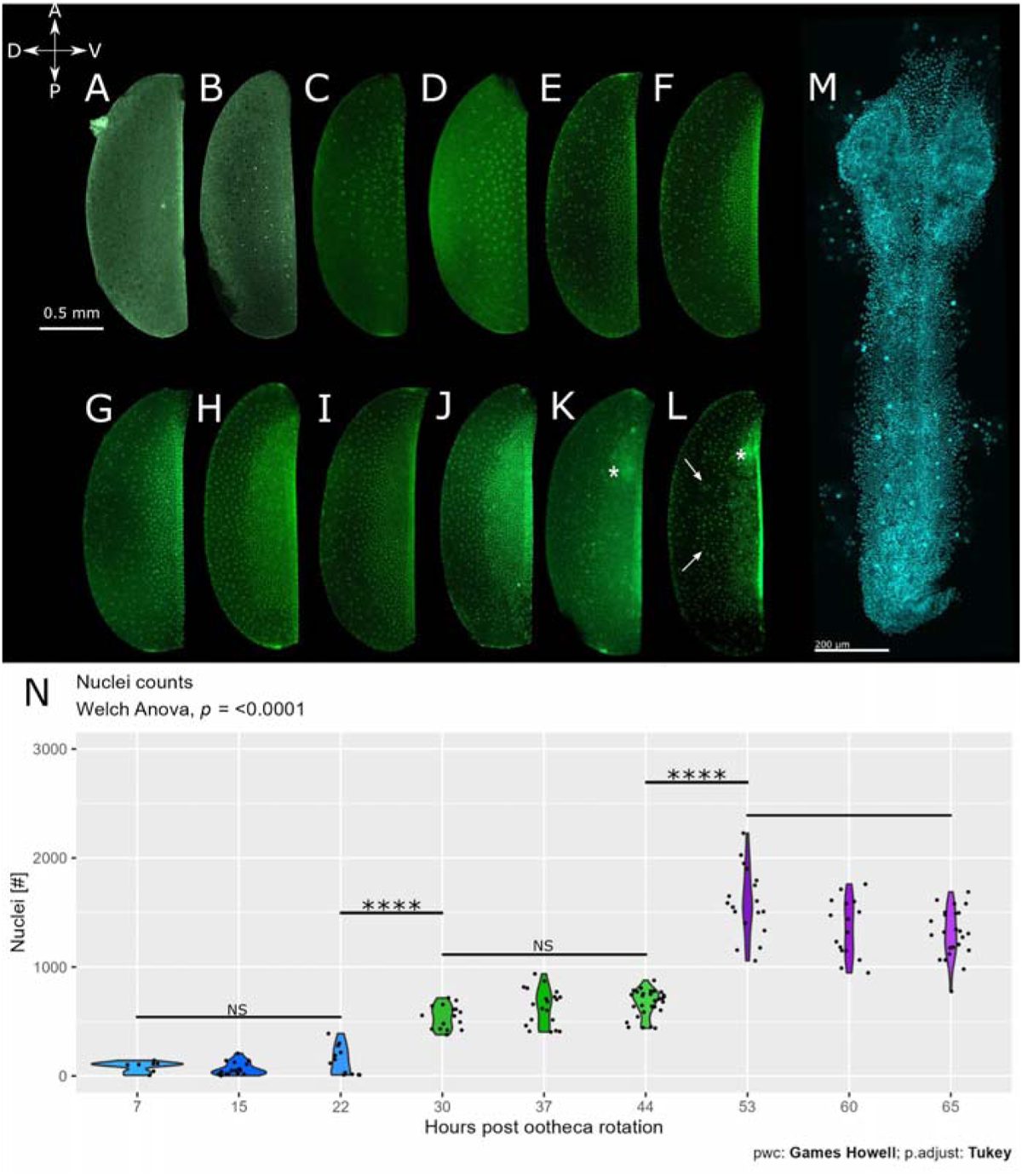
Nuclear dynamics in blastoderm and germband formation. Top panel: whole egg Sytox (A-L) and DAPI (M) staining. Embryos are arranged from youngest (A) to oldest (M). (A-L) Lateral view, ventral to the right, anterior top. (M) ventral view, anterior top, germband has been dissected out of the yolk. Asterisk in (K-L) marks the early head lobes, arrows mark the dorsal population of large nuclei. (N) Violin plot of nuclei counts taken at 7-8 hour intervals in the three days after ootheca rotation. Each black dot is the count of nuclei in an individual embryo. When the time points are divided into three bins (group A = 7,15,22 hr; group B= 30,37,44; group C = 53,60,65), there are significant differences in the number of nuclei in each bin (student’s t-test, p-value <2e-16).

*Stage 1. Days 0–1, 0-4% development (Fig. 1A-C): Cleavage and blastoderm formation stage.* This stage includes cleavage and blastoderm formation. After fertilization and fusion of the male and female pronuclei near the center of the egg (Fig. 1A), energids start to proliferate and migrate to the periphery of the egg. Nuclei first appear at the ventral surface of the egg, and later they appear at the dorsal edge (Fig. 1B-C). This stage continues until the embryo has ∼64-128 nuclei. At no stage do the energids show a uniform distribution over the surface of the egg – consistent with Anderson’s [5] description of a direct-differential blastoderm formation in Dictyoptera. The non-uniform distribution of energids across the surface of the egg could be explained by two phenomena: (1) non-random distribution of energid division, so that energids primarily divide on one half of the egg surface, or (2) energid migration, a scenario in which energid division occurs evenly across the surface of the egg, but the products of the division migrate to the ventral surface of the egg. Anti-phospho-histone 3 staining suggests the second explanation is correct during stage 1 and stage 2, as we observed dividing energids across the surface of the egg during these two stages (Fig. S1, n= 7 embryos in stage 1 and 11 embryos in stage 2).

*Stage 2. Days 1-2.5, 4-10% development (Fig. 1D-J): Syncytial blastoderm stage.* At this stage, the egg can be separated into 2 regions (Fig. 1D-J). The dorsal region is populated by a low density of energids, while on the ventral half of the egg, we find bilaterally paired regions of higher energid density. We hypothesize that energids from the dorsal region are migrating to the ventral region during this stage, but because of system limitations (i.e., no live imaging nor nuclear tagging and tracking), we cannot test this hypothesis explicitly. Cellularization occurs at the middle of this stage (see below, Cellularization section) (Fig. 2).

*Stage 3. Days 2.5-3.5, 10-14% development (Fig. 1K-L): Germ anlage/rudiment stage*. At this stage, two populations of nuclei can be identified in the dorsal half of the egg. One population consists of slightly larger, presumably polyploid [30–33] nuclei that are destined to become serosal cells (Fig. 1L marked with an arrow). The cells in the second population in the dorsal half of the egg are smaller and of equivalent size to the cells in the germ rudiment. We noticed a reduction in cell division on the dorsal side of the egg during stage 3 (Fig. S1, n=5 embryos counted.) As cell division continues on the ventral side of the egg, a subset of anterior cells further condense in the space where the head lobes will develop (Fig. 1K-L, marked with an asterisk).

*Stage 4. Days 3.5 and on (Fig. 1M): Early germband stage*. Following an accelerated phase of cellular condensation, the paired regions of higher cellular density fuse at the ventral region of the egg, starting at the posterior end and fusing towards the anterior, to form the bipartite germband (Fig. 1M). We define the end point of stage 4 as the complete fusion of lateral plates.

### Cellularization

We used wheat germ agglutinin (WGA) to look for the presence of cell membranes in embryos from oothecae 24-48 hours post extrusion. Using an Eclipse 80i Nikon Microscope, we detected evidence of cell membranes in only one of five embryos examined from an ootheca 24 hours post extrusion (Fig. S2). We took this embryo (Fig. 2B), plus a second embryo from the same ootheca with no discernable evidence of cell membranes (Fig. 2A), to an Olympus FV1200 confocal microscope for further imaging. In the embryo with evidence of cell membrane formation from the 24-hour-old ootheca, the WGA appeared to be condensing around nuclei (Fig. 2B’’). Surprisingly, the embryo without visible cell membranes had more nuclei than the embryo with membranes. The WGA signal is seen more tightly bound around nuclei in two embryos taken from an older ootheca between 24-40 hours post extrusion. By mid to late stage 2, embryonic nuclei are cellularized.

**Figure 2:**
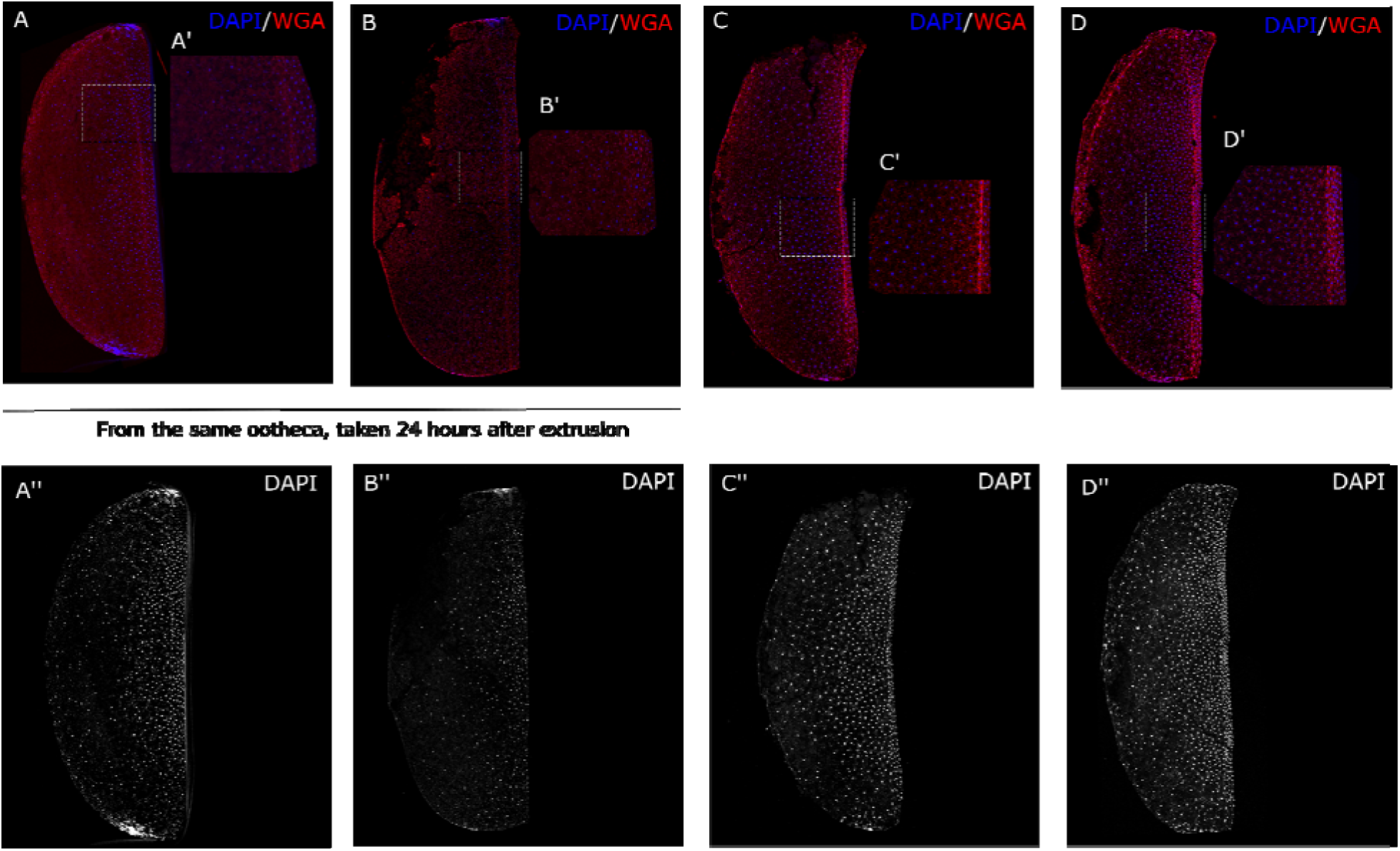
Cellularization occurs around stage 2. Embryos stained with DAPI and wheat germ agglutinin (WGA). Ventral side facing right. The top row shows both DAPI (blue) and WGA (red). White boxes show areas of higher magnification in panels A , B’, C’, and D’. The bottom row shows DAPI in grey from each embryo displayed in the panel directly above it. (A) Cell membranes do not appear visible with WGA in an embryo collected at 24 hours (4 percent of developmental time) after ootheca extrusion, but cell membranes are detected in embryo (B) collected from the same ootheca. (C, D) embryos collected from the same ootheca 24-48 hours (4-8 percent developmental time) post extrusion show visible cell membranes as detected with WGA.

### Segmentation

The determination of embryonic segments begins in embryonic stage 3 (Fig. 3). We used both chromogenic in-situ (cISH) and Hybridization Chain Reaction (HCR) to investigate the expression of *B. germanica hedgehog* (*Bg-hh*) (Fig. 3), an arthropod-wide marker of segment formation [34] and *B. germanica even-skipped* (*Bg-eve*) (Fig. 4), a gene that is higher in the segmentation cascade in insects, including *D. melanogaster* [35], *T. castaneum* [36], *N. vitripennis* [37], and *O. fasciatus* [38]. Lower imaging costs associated with cISH allowed us to obtain a time series tracking the appearance of each stripe of *Bg-hh* expression (Fig. 3), and HCR allowed us to observe the co-expression of both genes (Fig. 5). Shortly after head lobes become visible, *B. germanica hedgehog* (*Bg-hh*) expression appears (Fig. 3B, 5B). *Bg-hh* stripes appear one at a time, from anterior to posterior (Fig. 3B-H), although the timing of stripe appearance is not uniform. The antennal, mandibular, maxillary, labial, and first two thoracic segments appear rapidly and sequentially as the head lobes and germband condense (Fig. 5B-C). There is little detectable change in embryo morphology, as observed with DAPI stains, during the time these segments appear. Just before the two lateral plates of cells fuse in the head lobes, an eighth *Bg-hh* stripe (marking the third thoracic segment) appears (Fig. 5E). The embryo still shows eight *Bg-hh* stripes as the fusion process completes and the germband proper is formed (Fig. 5G).

**Figure 3:**
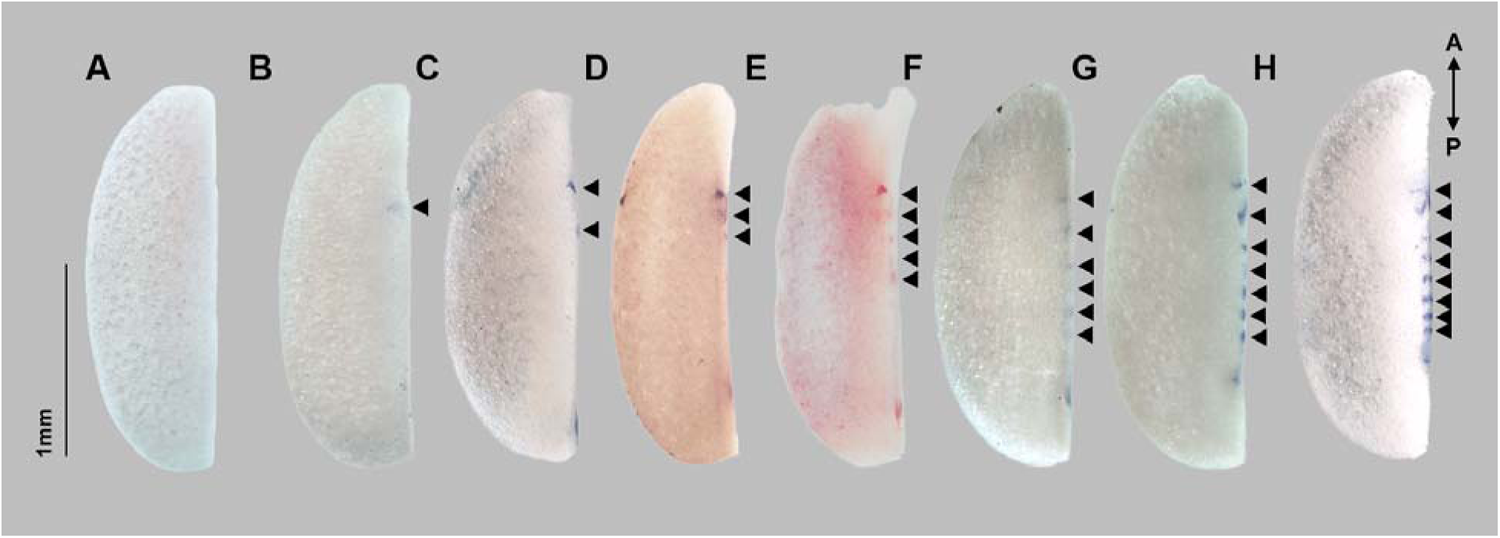
Sequential appearance of gnathal and thoracic *Bg-hh* stripes in *B. germanica*. Chromogenic in-situ for *Bg-hh*. The ventral side of the embryos faces right. Arrows indicate segments. Stripes of *Bg-hh* expression corresponding to each segment of the head and thorax appear individually.

We observed five *Bg-eve* stripes in an embryo shortly before *Bg-hh* appeared. These five stripes appear together in the middle of the anterior-posterior axis of the condensing germband, between the developing head lobes and what will become the segment addition zone (Fig. 4C-D, Fig. 5B-C). The five *Bg-eve* stripes persist as seven *Bg-hh* stripes appear, and they appear to mark the cells in the mandibular through the second thoracic segment. Note that pair-rule gene orthologs, such as *eve* are not known to be expressed in the pre-gnathal segments (ocular, antennal and intercalary) in any insects [38,39]. As the anterior-most *Bg-eve* stripe disappears, a new *Bg-eve* stripe appears posterior to the rest (Fig. 5D). This posterior-most stripe marks what will become the third and final thoracic segment. After *Bg-hh* expression appears in the third thoracic segment, we observed the number of *Bg-eve* stripes decline from six to three (Fig. 4D-G) as the number of *Bg-hh* stripes remains constant at eight (marking the ocular, mandibular and all gnathal and thoracic segments).

**Figure 4:**
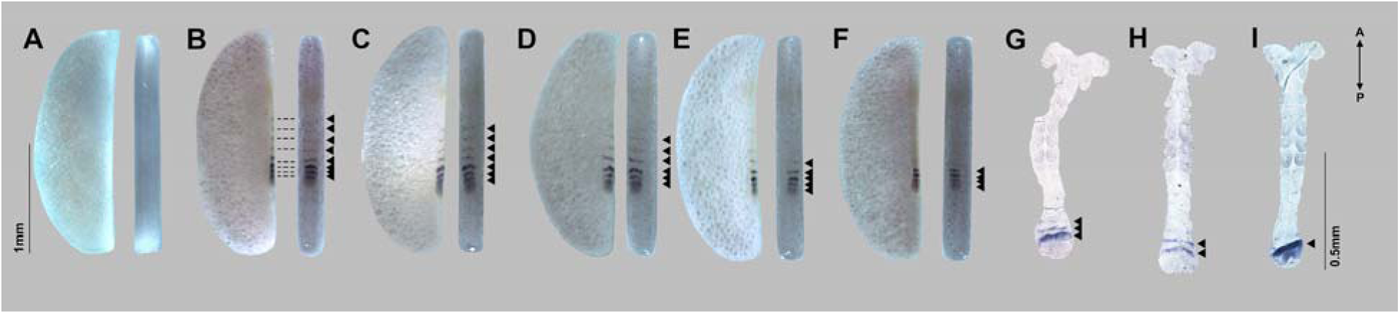
*Bg-eve* stripes appear simultaneously and disappear sequentially. Chromogenic in-situ of *even-skipped*. All embryos pictured are from the same ootheca. For each embryo, ventral view is shown on the left, lateral view is on the right, with the ventral side facing right. Dotted lines in B connect stripes of expression seen in the ventral and lateral views. Arrows highlight expression. Older embryos have fewer even-skipped stripes than younger embryos. *even-skipped* stripes associated with gnathal and thoracic segments are lighter in intensity than those associated with abdominal segments.

**Figure 5:**
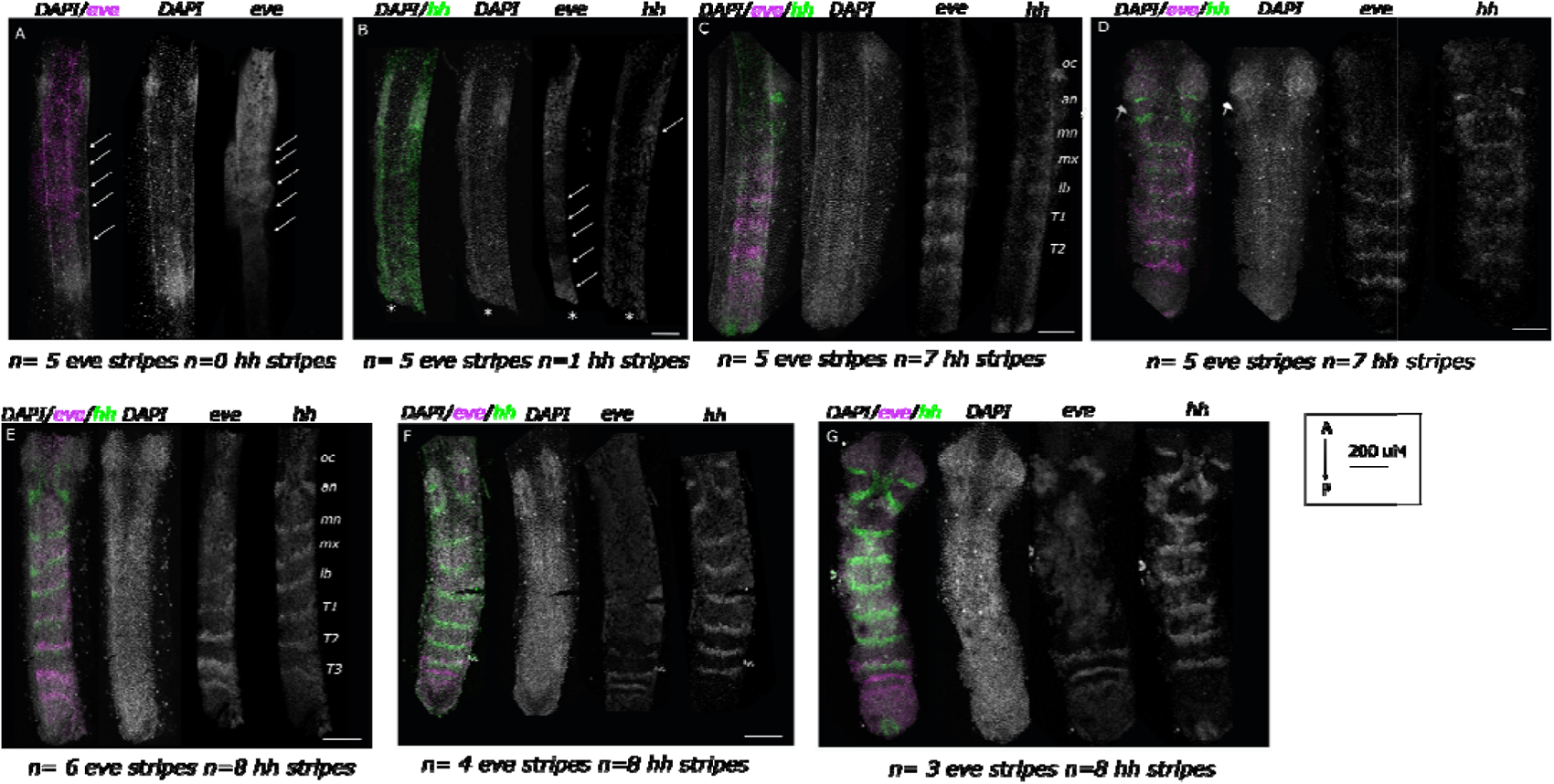
Gnathal-thoracic segmentation. Patterns of *Bg-hh* and *Bg-eve* were observed with HCR during gnathal thoracic segmentation (top panel). *Bg-eve* is shown in magenta, *Bg-hh* in green, and DAPI-stained nuclei in gray. Embryos are arranged from youngest (top left panel) to oldest (bottom right panel). The asterisk (*) in panel B indicates the embryo was broken at the posterior end. (A) *Bg-ev*e expression appears before *Bg-hh* during stage 3 of germband formation. We observed 5 stripes of *Bg-eve* expression (white arrows) at this point. No signal was detected for *Bg-hh*. (B) The first *Bg-hh* stripes to appear are in the head lobe. At this point, there are still five *Bg-eve* stripes, although the initial expression of *Bg-hh* appears uncorrelated with *Bg-eve*. (C and D) Antennal through the second thoracic segment appears rapidly in the condensing germband, indicated by the jump from one to seven *Bg-hh* stripes with little to no change in embryo morphology. (E) The number of *Bg-eve* stripes increases from five to six as the number of *Bg-hh* stripes increases from seven to eight. (F, G) *Bg-eve* stripes gradually disappear as the number of *Bg-hh* stripes remains constant.

During abdominal segmentation, cells first express *Bg-eve* before expressing *Bg-hh*. *Bg-eve* expression fades gradually in each segment as the *Bg-hh* expression strengthens (Fig. 6). *Bg-eve* stripes appear one by one from the posterior as they fade anteriorly, leaving a more or less constant number of three stripes in the anterior segment addition zone.

**Figure 6:**
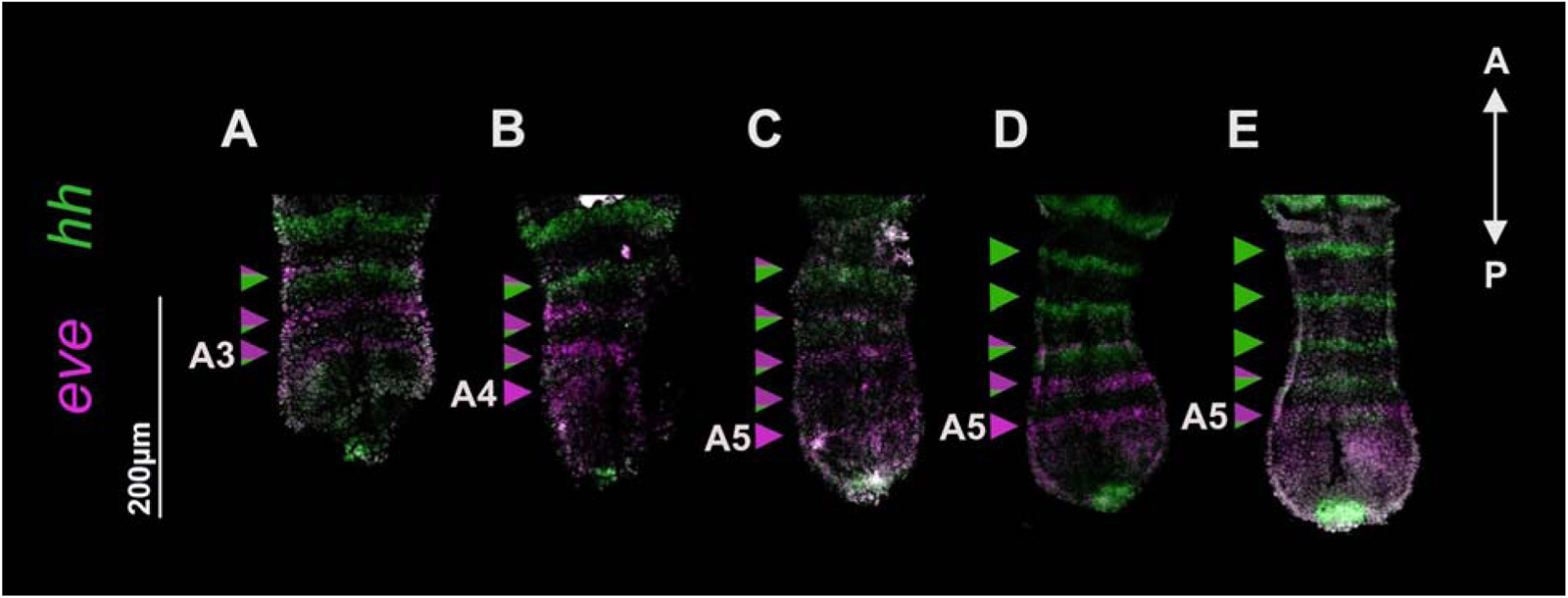
Abdominal segmentation: During abdominal segmentation, *Bg-eve stripes* disappear as *Bg-hh* stripes appear. *Bg-eve* is shown in magenta and *Bg-hh* in green. Embryos are arranged from youngest (left) to oldest (right). Arrows to the left of the embryos highlight each stripe with a proportional representation of green or magenta to indicate how many cells in the stripe are expressing *Bg-eve* (magenta) versus *Bg-hh* (green). A3 = abdominal segment 3, A4=abdominal segment 4, A5 = abdominal segment 5.

### Axial patterning

The homeobox transcription factor and posterior determinant *caudal* (*cad*), and the zinc-finger transcription factor and anterior determinant *hunchback* (*hb*) were used to observe the process of early axial patterning (i.e., determination of anterior and posterior of the embryo). Both genes are conserved blastoderm patterning genes in arthropods and Bilateria generally [34,41]. In *B. germanica* embryos, cISH and HCR showed no *Bg-cad* or *Bg-hb* expression in the early blastoderm embryo (data not shown). This finding is corroborated by qPCR experiments (Fig. 7A), and suggests that *Bg-cad* and *Bg-hb* mRNA are not maternally deposited. Zygotic transcription of *Bg-hb* is initiated 2 days (8 percent development) post ootheca extrusion. We observed a local peak of *Bg-hb* expression at 3 days (12 percent development) post ootheca extrusion. *Bg-cad* expression initiates 3 days (12 percent development) post ootheca extrusion and maintains its expression level throughout day 4. In the early germband stage of embryogenesis, HCR shows *Bg-cad* is expressed at the posterior end of the embryo (Fig. 7B). This broad domain of expression remains as the germband continues to elongate (Fig. 7B). Using HCR, no clear *Bg-hb* expression was observed at the early germband stage (Fig. 7B). Later, as the germband elongates, *Bg-hb* is expressed in paired lateral cells along the midline, and in a branching pattern into the head lobes.

**Figure 7:**
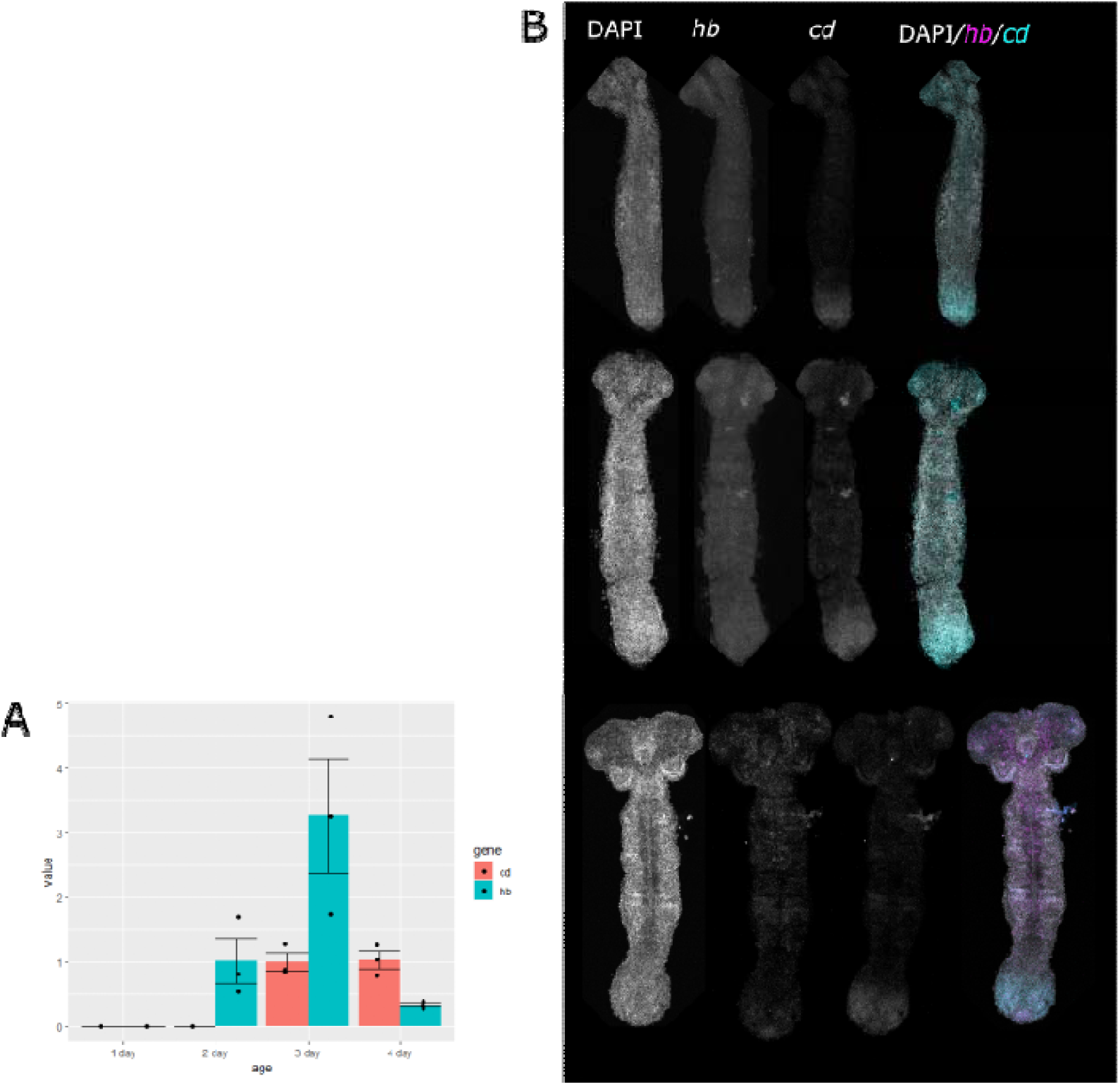
Early expression of *hunchback* and *caudal*. (A) qPCR measuring *Bg-hb* and *Bg-cad* expression relative to actin levels in embryos from oothecae one, two, three, and four days post extrusion. *Bg-hb* expression peaks at 3 days (12 percent development) post ootheca extrusion. (B) HCR of both genes in embryos three days post extrusion. Embryos in top and middle row are approximately 4 days post extrusion. Embryo in the bottom row shows neuronal expression of *Bg-hb* and is approximately 5 days post extrusion.

### Mesoderm formation

The bHLH encoding gene *twist* (*twi*) is a conserved myogenic and mesoderm marker within arthropods[21,42–46] and in other bilaterians [47–50]. In insects, it also has a central role in the dorso-ventral patterning pathway [51–53]. We investigated the expression of *Bg-twi* from the blastoderm through to the germband stage in order to follow the initial stages of mesoderm formation. During the condensation of the germ anlage at the beginning of stage 3, a punctate pattern of expression can be seen on the posterior-ventral region (*Fig. 8A-C)*. As the headlobes fuse in the germband (stage 4), *Bg-twi* is visible in the posterior germband and in the lateral head lobes, gradually expanding to cover almost the entire embryo (*Fig. 8D-I)*. During posterior segmentation, *Bg-twi* expression remains in the limb-buds only, while a new expression pattern appears as segmental stripes in recently formed segments (*Fig. 8J)*.

**Figure 8:**
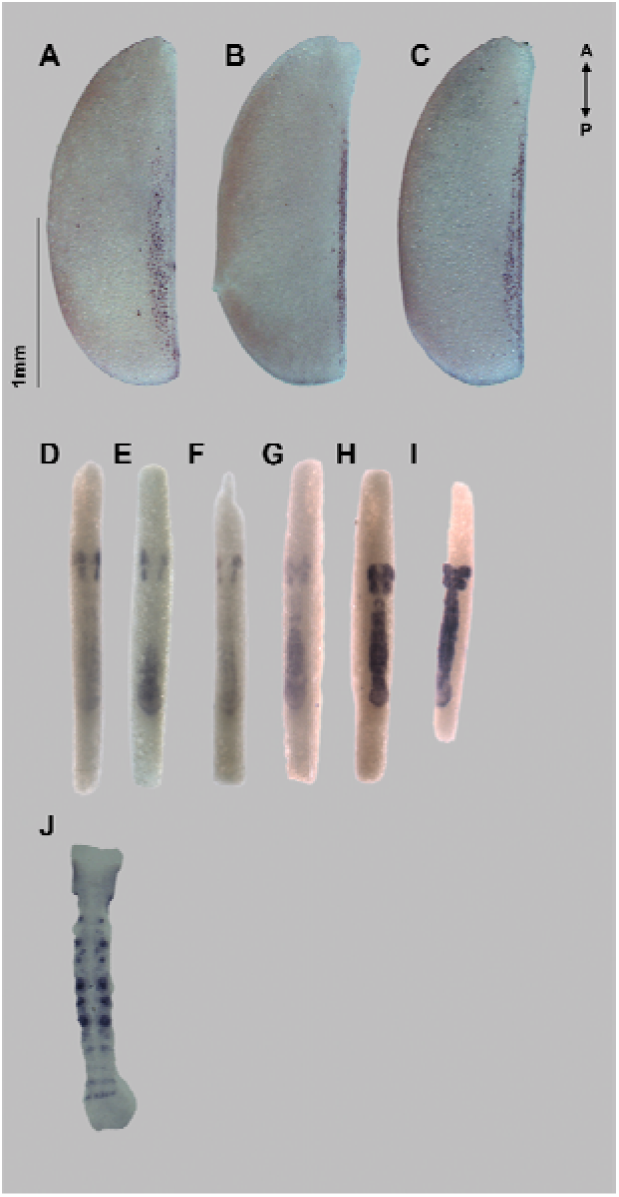
Mesoderm formation. chromogenic in-situ of *Bg-twi*. All embryos in a row are from the same ootheca. (A-D) Germ rudiment (stage 3), embryos from lateral view are presented with the ventral side to the right. Individual cells marked as mesoderm are visible in a ventral position. Considering the unusual shape of the egg this is consistent with the classical expression pattern as seen in *D. melanogaster* (Leptin, 1991) and *T. castaneum* (Handel et al., 2005). (E-H) germband fusion through early germbands (stage 4 and on) ventral view of hacked eggs. The expression can be seen from the earliest stages in the presumptive head lobes and the posterior of the forming germband. The expression advances anteriorly with the fusion of the embryo proper. A clear patch is maintained for some time between the PGS and gnathal segments. (M) As limb primordia begin to form, *Bg-twi* becomes segmental and marks the formation of segmental mesoderm.

## Discussion

We investigated the early stages in the embryonic development of the German cockroach, *Blattella germanica*, as an emerging model for evo-devo studies. *B. germanica* is a member of a basally branching taxon relative to the well-studied Holometabola, and its phylogenetic position in an underrepresented clade and in relation to other species studied can provide a key reference point for developmental studies. In this work, we have described some of the most fundamental developmental events in the earliest stages of development, including blastoderm and germband formation, cellularization, mesoderm formation, embryo polarization, cellularization, and segmentation. We hope that these data will provide a developmental roadmap for *B. germanica,* highlighting unique aspects of its embryogenesis that can be compared to other insect species. Furthermore, we hope this study will stimulate additional research utilizing this emerging model organism.

### The development of *Blattella germanica*

Early embryogenesis can be divided into four stages. Stage 1 involves cleavage and formation of a direct-differentiated blastoderm with nuclei migrating from the ventral to the dorsal surface. In stage 2, the syncytial blastoderm exhibits higher nuclear density ventrally compared to dorsally, suggesting nuclear migration. Cellularization occurs midway through this stage. Stage 3 sees the formation of a germ anlage/rudiment with larger polyploid nuclei dorsally, giving rise to serosa. Meanwhile, ventrally, the germ rudiment nuclei condense anteriorly, where head lobes will form. In stage 4, the germ rudiment’s lateral plates fuse ventrally from posterior to anterior, forming the bipartite germband.

The anterior determinant *Bg-hb* and the posterior determinant *Bg-cad* are not expressed in early developmental stages, and only come up in the germband. This is contrary to what we had expected based on data from other species. However, in the milkweed bug *Oncopeltus fasciatus*, as well as in the pea aphid *Acyrthosiphon pisum* (both members of Hemiptera)*, cad* is not expressed maternally [54–56]. Similarly, *hb* was not found to be significantly expressed in early stages of *O. fasciatus* [55]. Combined with our results in *B. germanica,* this suggests that in hemimetabolous insects in general, and possibly ancestrally for insects, early polarization may not be driven by maternal transcription factors. Instead, it may be driven by structural elements related to the morphology of the egg.

We have not followed gastrulation and mesoderm formation in sufficient detail to draw strong conclusions about how they occur in *B. germanica*. However, the expression pattern of *Bg-twi* we do see is quite similar to that reported for the holometabolous beetle *Tribolium castaneum* [21].

Segmentation in *B. germanica* follows the intermediate-germ paradigm. The determination of embryonic segments begins in stage 3, marked by the almost simultaneous segmentation of the pre-gnathal and gnatho-thoracic segments. During abdominal segmentation, *Bg-eve* precedes *Bg-hh* in each segment before fading as *Bg-hh* expression strengthens.

### Evolutionary implications

Researchers in the early 20th century (reviewed by [5,18,19]) discussed the formation of the insect blastoderm and noted two distinct phenomena during this phase. The first is the emergence pattern of nuclei on the egg’s surface. This emergence can be uniform across the whole egg, as in *Oncopeltus fasciatus* (manuscript in preparation), or more concentrated on a specific region, usually the postero-ventral region, as in *B. germanica*. The second phenomenon is the final distribution of nuclei on the egg’s surface. Following a local emergence, nuclei migrate across the surface and are eventually distributed uniformly, as in the cricket, *Gryllus bimaculatus* [57], or remain localized, as in *B. germanica*. Indeed, recent work has shown that even among species whose embryos all possess a uniform blastoderm phase, different processes control cell migration and nuclear division in this early stage [58]. Little work, though, has investigated the relationship between variation in blastoderm type—uniform versus direct-differentiated—and subsequent early germband patterning. For example, embryos with direct-differentiating blastoderms lose synchronous cleavage earlier than those with uniform blastoderms [5]. We found that the direct differentiating blastoderm of *B. germanica* does not affect the expression patterns of the genes we studied. Most of the genes investigated in this paper were expressed in patterns similar to those previously reported in insects with uniform blastoderms. Preliminary results from a literature review (manuscript in preparation) suggest that the direct-differential blastoderm (regionalized emergence followed by regionalized final distribution) is an apomorphy of Polyneoptera.

### Segmentation and tagmatization

The timing and dynamics of the segmentation process are summarized in Fig. 9. We have shown a transition in segmentation mode between the thoracic and abdominal segments during the development of *B. germanica*. At the level of segment-polarity genes, the process seems to be sequential throughout but more rapid in the early stages of segmentation. Looking at the expression of *Bg-eve* (a pair-rule gene in *Drosophila melanogaster*) representing a higher regulatory tier [34], it appears that the gnathal and thoracic segments exhibit a more-or-less simultaneous early determination phase, similar to the progressive segmentation described for the wasp *Nasonia vitripennis* [59], while the abdominal segments are patterned via a classical segment addition zone (SAZ) with a cycling process of segmentation, as shown in the holometabolous *T. castaneum* [60,61], the hemimetabolous *O. fasciatus* [29,62] and several non-insect arthropods [63–65]. The distinction between the two segmentation modes is not as sharp as in *O. fasciatus*, where there is a transition from a blastoderm to an internalized germband between the two. However, gnatho-thoracic segmentation in *B. germanica* occurs within the early determined embryonic rudiment, whereas abdominal segmentation occurs from a posterior SAZ that is only determined towards the end of thoracic segmentation. The third thoracic segment is unusual in exhibiting a transitional mode. It is patterned later than the other thoracic segments but is still distinct from the SAZ-mediated patterning of the abdominal segments. This transition in segmentation dynamics between the thorax and the abdomen, similar to that reported in O. fasciatus, suggests that an early developmental boundary between these tagmata may be a more broadly conserved feature than previously recognized.

**Figure 9:**
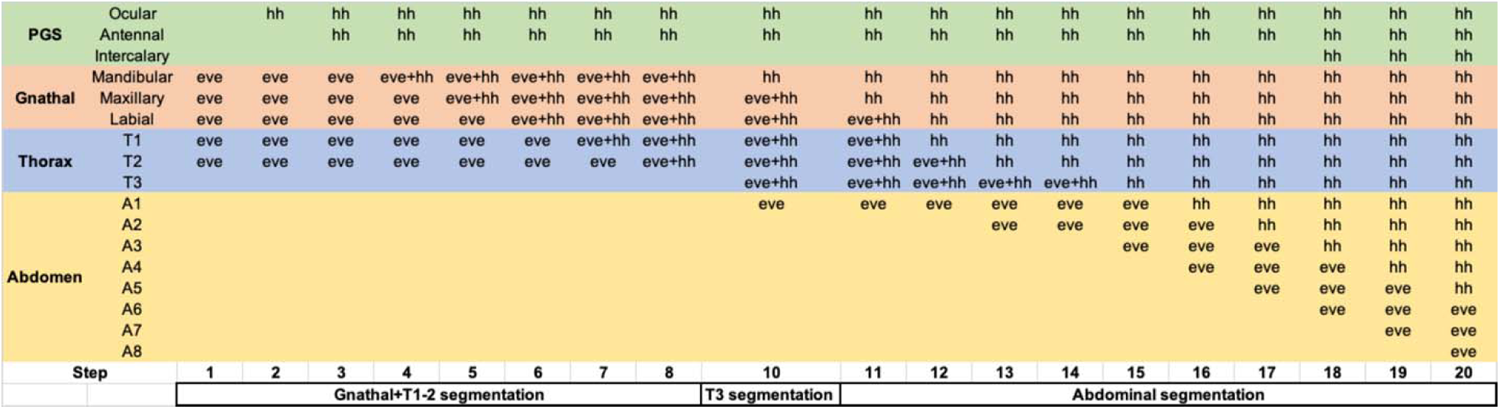
Segmentation Dynamics. Summary of *Bg-eve* and *Bg-hh* expression data over time. Each row represents a segment, with the columns representing successive time-points (steps) in the developmental process. The steps are arbitrary points in which there is a change in expression pattern relative to the previous step, and are not to scale. Data accumulated from numerous single- and double-stained embryos, not all of which are shown in the Results.

### Follow up experiments

We presented a rough time course of Bg-twi expression in the B. germanica embryo, but believe a more detailed time series would be valuable. Specifically, high resolution images combining Bg-twi probes with DAPI could illuminate how mesoderm formation proceeds as the lateral plates of the B. germanica embryo come together. Because the lateral plate model is a suggested apomorphy of Polyneoptera, understanding mesoderm formation in this type of embryo would be extremely valuable.

## Materials and Methods

### Insect husbandry

The *B. germanica* colony was grown from several starting colonies supplied by Meital Labs (https://www.meital-labs.co.il/). Insects were kept at 25℃ and ∼40% humidity with a 14:10 D:L cycle (Humidity and temperature values fluctuate by ∼10% on a seasonal basis). Insects were reared in large containers with perforated covers. Each container is 19x19x22 cm, with several dozen individuals of continuous generations. In the container, a few cardboard tubes were used as shelter. Insects were fed dog chow and dried oats *ad libitum*. Distilled water was provided in 50 ml Erlenmeyer flasks sealed with water-soaked cotton balls. The cages were cleaned, and the food and water were renewed weekly.

### Embryo staging

As oothecae emerge from female *B. germanica*, the seam of the egg case is oriented dorsally (Fig. S3A). After all eggs are laid and the ootheca is fully extruded, it rotates 90 degrees, such that its seam is aligned laterally (Fig. S3B). We collected adult females laying eggs (identified by a partial extruded egg case), and we also collected females with oothecae that had extruded but were not rotated. A fully extruded but not rotated ootheca (Figure 1D in ref. [3]) was set as time point 0. Since the oothecae are not viable after being removed from the mother, collected females were isolated until their embryos had reached the desired age, at which point we manually removed the oothecae for further study.

### Egg case removal and fixation

Staged females were anesthetized with CO2 or cooled on ice before manual removal of oothecae. Oothecae were placed into 1 mL of water, then heated for 10 minutes at 85℃ and subsequently submerged in ice for up to five minutes. Except for embryos used for wheat germ agglutinin (WGA) staining, embryos were dissected out of the ootheca in phosphate-buffered saline with Tween (PBST) and placed directly into fixative (4% formaldehyde in phosphate-buffered saline). Embryos were fixed in 4% formaldehyde for one hour and then gradually dehydrated to 100 percent methanol. Embryos were stored in methanol at -20°C for at least overnight, up to several months. For some nuclear stains and immunohistochemistry, the methanol step was skipped. For embryos used in WGA staining, dissection was done in phosphate-buffered saline with no detergent, and fixation was performed for twenty minutes in 4% paraformaldehyde in phosphate-buffered saline.

### Nuclear staining

If necessary, embryos were rehydrated gradually from methanol and washed for an additional 5 minutes in PBST. Fixed embryos were stained with 1 µL Sytox green (Invitrogen, Sytox Green Nucleic acid, S7020) in 1 mL PBST and incubated at RT, in the dark, for 20 minutes. Stained embryos were washed twice in the dark for three minutes with PBST solution. When staining with DAPI, we added 0.5 µL DAPI (Invitrogen, DAPI, D1306) in 1mL PBST and incubated at RT in the dark for 15 minutes. Stained embryos were washed once for 15 minutes with PBST in the dark.

### Immuno-fluorescence

Before adding primary antibodies, the embryos were incubated in a blocking solution (1% BSA (Bovine serum Albumin, MP Biomedicals cat no. 160069) and 5% NGS (Normal Goat Serum, Vector Labs cat no. S-1000) in PBST) for one hour. Embryos were incubated with anti-phosphorylated histone H3 (PH3) antibody (1:500; Abcam, ab14955) or Anti-alpha tubulin (1:500; DSHB,12G10) overnight. The following morning, embryos were washed four times for 15 minutes with PBST. Secondary antibodies (1:200; Alexa fluor 448/546/594, anti-mouse, Invitrogen) were added to the blocking solution, and embryos were incubated in the dark for 2 hours.

#### Gene sequence isolation

All gene sequences were identified in tBLASTn searches of *B. germanica* gene models on the i5k database (bger_OGS_v1.2, url: https://i5k.nal.usda.gov/data/Arthropoda/blager-%28Blattella_germanica%29/Bger_2.0/2.Official%20or%20Primary%20Gene%20Set/). *Bg-hh* (BGER011018-RA-CDS) was identified after a tBLASTn search using hedgehog protein from *Oncopeltus fasciatus* (NCBI accession number AYR04649.1) as bait. *Bg-hb* (gene model BGER015825-RA-CDS) was identified in a tBLASTn search using the *hunchback* protein from *D. melanogaster* as bait (Fly Base FBgn0001180). *Bg-cad* (gene model BGER016043-RA-CDS was identified using caudal protein from *D. melanogaster* as bait (Fly Base FBgn0000251). *Bg-twi* (gene model BGER025575-RA-CDS) was identified using *D. melanogaster* twist protein (Fly Base FBgn0003900) as bait. *Bg-eve* was identified using an *even-skipped* sequence from *Drosophila melanogaster* as bait (Fly Base FBgn0000606). The *B. germanica* gene model retrieved as a top hit in a tBLASTn search for *eve* (BGER013241-RA-CDS) was only 166 amino acids. When aligned to other insect even-skipped proteins, this *B. germanica* gene model was missing sequence from the C-terminal end. We then used a sequence from the *Cryptotermes secundus even-skipped* protein (XP_023719287.1) as bait in a tBLASTn search of the *B. germanica* genome in an attempt to retrieve sequence from exons downstream of the *B. germanica* gene model with the putative *eve* sequence. This strategy was successful; we obtained what appeared to be a 317 bp exon downstream of the original *B. germanica* gene model. Using this exon as a template for a reverse primer and the gene model (BGER013241-RA-CD) as a template for a forward primer, we designed primers that then amplified a 916 bp fragment of *B. germanica even-skipped* from embryonic cDNA. This sequence was deposited in NCBI with the accession number (TBA).

### Chromogenic in-situ hybridization probe preparation

We collected total RNA from three-day-old oothecae manually removed from the female. RNA was extracted with Trizol (Invitrogen cat no. 15596026) using the manufacturer’s protocol. One microgram of RNA was used in a reverse transcription reaction primed with a 1:1 mixture of oligo dT:random hexamers. We used Bioline’s RTase (cat No BiO-27036). Templates for *Bg-hh*, *Bg-eve, Bg-hb*, *Bg-cad,* and *Bg-twi* probes were amplified using Tiger polymerase from Hylabs (EZ-2031), with primers that had a T7 sequence added to the 5’ end of the reverse primer. To amplify templates for the *Bg-hh* probe, we used the following cycling parameters: 95°C for 3 minutes, followed by 32 cycles of 95°C for 30 seconds, 55°C for 30 seconds, and 72°C for 30 seconds, with a 3-minute extension at 72°C at the end of the cycles. Other probes were amplified with the same cycling parameters but different annealing temperatures (see PCR conditions table) Templates were purified using Macherey-Nagel’s NucleoSpin Gel and PCR Clean-up kit (740609.50). We synthesized DIG-labelled probes with T7 polymerase from Roche (10881767001).

### Chromogenic In-situ Hybridization

Embryos were gradually rehydrated from 100 percent methanol. After 3 × 1-minute PBST washes, a post-fixation of 20 min in 4 percent formaldehyde in PBST and another round of 3 × 1-minute PBST washes were carried out. Embryos were incubated in a hybridization buffer at 65 °C for 5 min, then the hybridization buffer was refreshed, and embryos were incubated again in a hybridization buffer at 65 °C for 2–4 h. DIG-labeled probes were applied at a 1 ng/uL concentration in the hybridization buffer and left overnight at 65 °C. The following day, the following washes were performed: 20 min in preheated hyb buffer (65 °C); 3 × 20 min in 2x SSC/0.1% Tween-20 (65 °C); 3 × 20 min in 0.2x SSC/0.1% Tween-20 (65 °C); 2 × 10 min in PBST at room temp; 1 h in blocking solution (5 percent sheep or goat serum, 2 mg/mL BSA, 1 percent DMSO in PBST); 4 h in 1:1500 solution of anti-digoxigenin antibody (Roche, cat no. 11333089001):blocking solution; 3 × 1-minute PBST washes, and left overnight in PBST at 4 °C. The following day, embryos were washed in the following series: 4 × 20 min wash in PBST; 2 × 10 min of washing in freshly prepared staining solution (0.1 M 9.5 TrisHCl; 0.05M MgCl2; 0.1M NaCl; 0.1M Tween-20); 1 × 10 min of staining solution + PVA (0.1 M 9.5 TrisHCl; 0.05M MgCl2; 0.1M NaCl; 0.1M Tween-20; 0.025 percent polyvinyl alcohol). 20 μL of NBT/BCIP from Roche in 980 μL of staining solution + PVA was used for color development, which continued up to 4 hr. Staining reactions were stopped when we observed background stain developing. Staining reactions were stopped with PBST washes and fixation in 50 percent methanol:PBST. Embryos were imaged in 70 percent glycerol.

### Hybridization chain reaction

We used probes designed by Molecular Instruments for *Bg-cad* (lot number PRP735), *Bg-hb* (lot number PRP734), *Bg-hh* (lot number PRK420), *Bg-eve* (lot number PRK419), and *Bg-twi* (lot number RTG920). For probe design, we provided Molecular Instruments with the entirety of each gene’s available coding sequence. Probes designed by Molecular Instruments target several dozen nucleotides each and span the entirety of the coding sequence in a non-contiguous fashion. Probes for *Bg-hb* and *Bg-hh* were all designed to anchor an amplification reaction with a 488 fluorophore. Probes for *Bg-eve*, *Bg-cad*, and *Bg-twi* were designed to anchor an amplification reaction with a 594 fluorophore.

For HCR-RNA FISH, a modified version of the protocol described by Bruce et al. [66] was used. Hybridization buffer, wash buffer, and amplification buffer were provided by Molecular Instruments, while we made the detergent solution (1 percent SDS, 0.5 percent Tween, 50 mM Tris-HCl (pH 7.5), 1 mM EDTA (pH 8.0), 150 mM NaCl) as described by Bruce et al. [66]. Embryos were rehydrated from methanol to PBST as described above in the chromogenic in-situ section and then washed 1 × 10 min and 2 × 5 min in PBST. Embryos were incubated at RT for 30 min in the detergent solution, then 30 min at 37 °C in 200 μL of hybridization buffer. Probes were prepared at a concentration of approximately 10 nM, made by diluting 1.6 μL of the 1 μM stock from Molecular Instruments into 150 μL of hybridization buffer. Embryos were incubated overnight at 37 °C in the probe solution. The following day, embryos were washed in 1 mL of wash buffer at 37 °C (4 × 15 min), followed by 2 × 5-minute washes at RT with 5% SSCT (5x sodium chloride sodium citrate with 1 percent Tween). Embryos were incubated in an amplification buffer (Molecular Instruments) for 30 min at RT, while hairpins were prepared by heating to 95 °C for 30 seconds, followed by 30 min at RT in the dark. 4 μL of each hairpin were added to 100 μL of the amplification buffer. This solution was applied to embryos after removing the 1 mL of the amplification buffer used for the pre-amplification step. Embryos were then incubated overnight in the dark at RT. Embryos were washed the next morning with five percent SSCT washes at RT (volume 1 mL) twice for 5 min and once for 15 min. We then did a 15-minute wash with 1 μL DAPI into 1 mL of five percent SSCT, followed by two 15-minute washes with five percent SSCT. Embryos were placed in 50% glycerol/PBS for 30 minutes before being transferred to 70% glycerol in PBS and mounted on slides.

### Wheat germ agglutinin staining

Immediately after a twenty-minute fix in 4% paraformaldehyde, embryos were washed for 3 x 5 minutes in phosphate-buffered saline (PBS). Embryos were then transferred to WGA (Invitrogen, Wheat Germ Agglutinin (WGA), W21405) in PBS (1 ng/mL) and incubated for thirty minutes. Embryos were then washed for 3 x 10 minutes in phosphate-buffered saline, with the second wash containing DAPI at a concentration of 5 ng/mL. Embryos were mounted and visualized in seventy percent glycerol.

### qPCR

Embryos were boiled and dissected in PBST as described above: “Egg case removal and fixation.” Upon dissection, Embryos were immediately placed in Trizol. A single sample contained all embryos from one ootheca. RNA was extracted according to the manufacturer’s protocol. Two hundred to three hundred nanograms of RNA were put into a reverse transcription reaction. Watchmaker StellarScript reverse transcriptase was used to generate cDNA, with the following parameters used in the thermocycler: 25℃ for 10 minutes, 42℃ for 20 minutes, and then 80℃ for 10 minutes. qPCR primers were designed to span exon junctions or in two separate exons. Primer efficiency was validated with a dilution series. qPCR was performed on QuantStudio3 from ThermoFisher using Fast SYBR Green Master Mix (cat no 4385610) in 10 uL reactions. All reactions were performed using technical triplicates. Each biological replicate contained all the RNA from a single ootheca. We ran *cyclophilin* and *actin* as internal controls for each biological sample; however, we could not amplify cyclophilin in one-and two-day-old oothecae. We used the Pfaffl method to normalize expression values of *caudal* and *hunchback* to the geometric mean of *actin* and *cyclophilin* expression for samples from three and four-day-old oothecae. We then re-ran the analysis, normalizing *caudal* and *hunchback* expressions to *actin* alone. Results did not change considerably when expression of *caudal* and *hunchback* were normalized to *actin* alone, so we proceeded to use only *actin* for further analysis so that three and four-day samples could be compared to samples from one and two-day-old oothecae.

### Imaging

Egg case and embryo dissections were done under a 2000-Stem ZEISS dissecting scope.

Images of ISH and IF stained embryos were captured using a Nikon ‘digital sight’ console connected to a DS-Fi1 digital camera mounted on either an SMZ1500 Nikon dissecting scope or an AZ100 zoom stereoscope. Images of slide-mounted embryos were captured with the same console and camera mounted on an Eclipse 80i Nikon Microscope.

HCR images were acquired with an Olympus FV1200 confocal based on an IX-83 inverted microscope (Olympus, Japan), using a 10x/0.4 or 40×/0.95 air objective. Confocal fluorescence images of DAPI, Alexa 488 anti-PH3, Alexa 488 anti-alpha tubulin, and DIC images were acquired. The DAPI channel used 405 nm excitation and a 430-470 nm emission, and the Alexa 488 channel used 488 nm excitation and a 500-540 nm emission, acquired sequentially. Z-Z-stacks were acquired with 1.5 or 4 μm stepping.

Images were processed in Fiji (Fiji is just ImageJ) [67]. We selected and projected the Z-stacks containing the signal for each fluorescent channel, as determined subjectively by the eye. After projecting stacks, we adjusted the contrast and brightness of each channel (using the window, level, contrast, and brightness scales in Fiji) to maximize the signal from the HCR. We then false-colored and merged channels.

## Supplementary files

**Supplemental Figure 1:**
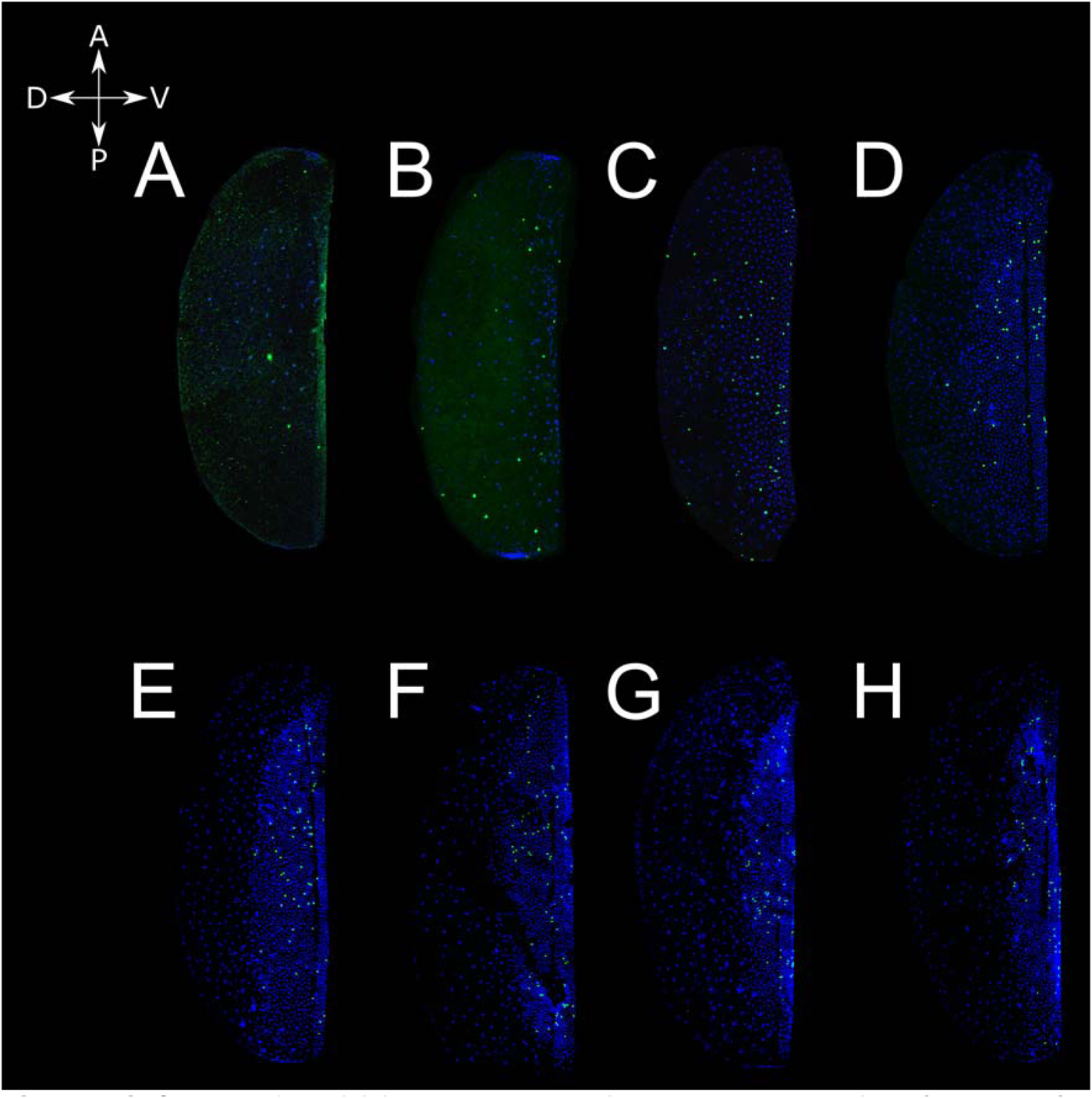
Cell division becomes restricted to the ventral side of the egg after the formation of the germ rudiment. DAPI (blue) and anti-phospho-histone H3 (green) mark nuclei and dividing cells in embryos stage 1-4. Cell division is observed across the surface of the egg in embryonic stages 1 and 2 (panels A-C), but once the germ rudiment forms, cell division occurs overwhelmingly in the area of the forming germband (panels D-H).

**Supplemental Figure 2:**
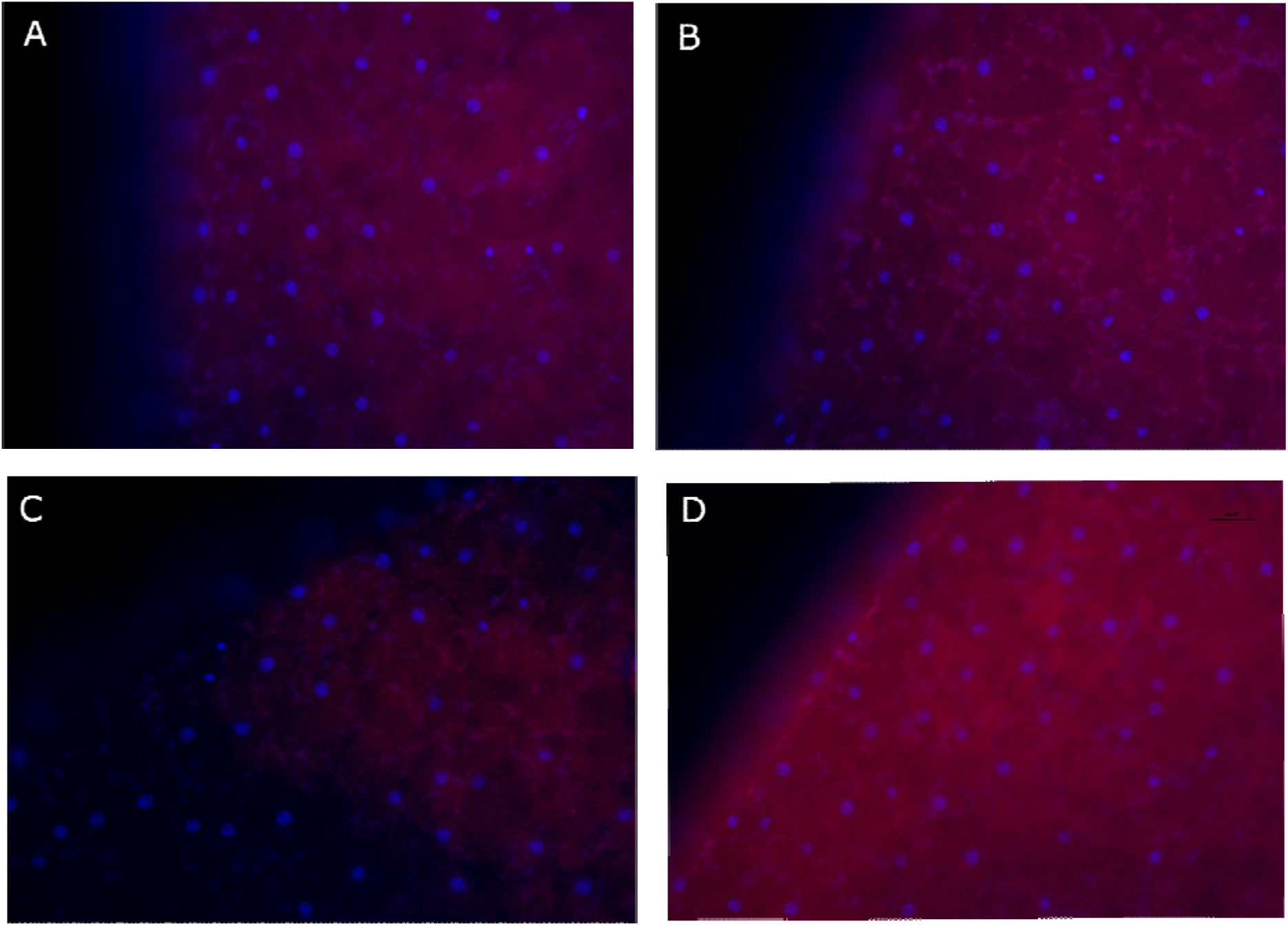
Representative images from 4 different embryos from the same egg case taken 24 hours post extrusion (4 percent of developmental time). Only the embryo in B shows a discernable signal of WGA staining around nuclei which is distinct from the background.

**Supplemental Figure 3:**
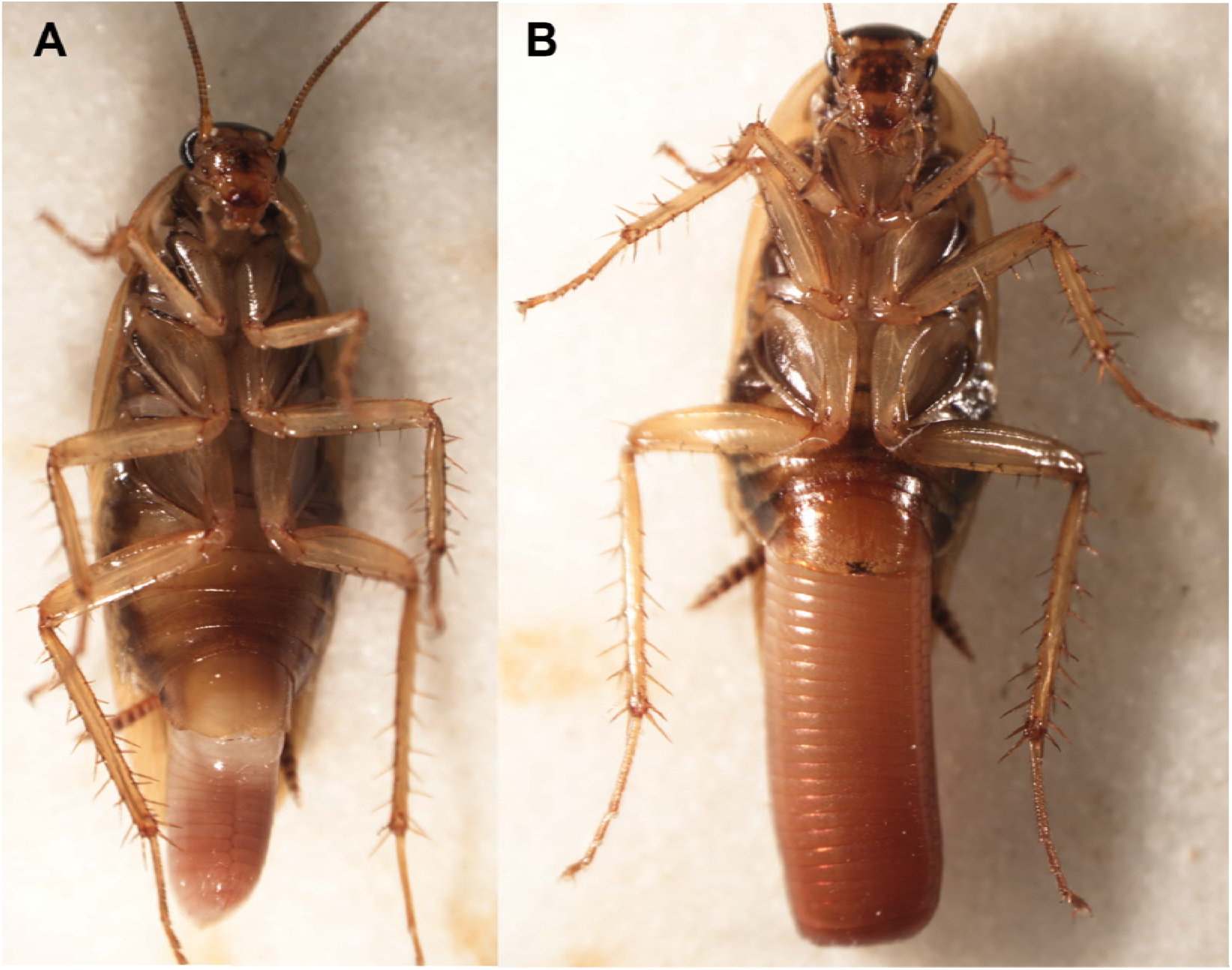
Ootheca protrusion in the German Cockroach female. As the egg case protrudes from the posterior abdomen of the female, the seam of the egg case is oriented dorsally. Eggs are laid successively into individual compartments After all eggs are laid and the ootheca is fully extruded, it rotates 90 degrees, such that its seam is aligned laterally.

**Supplementary table 1:**
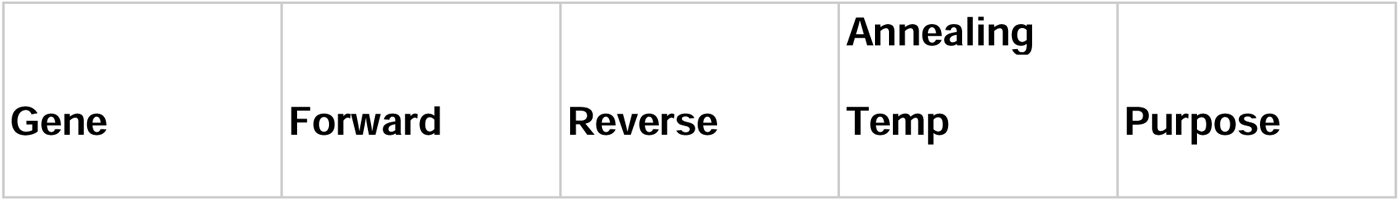
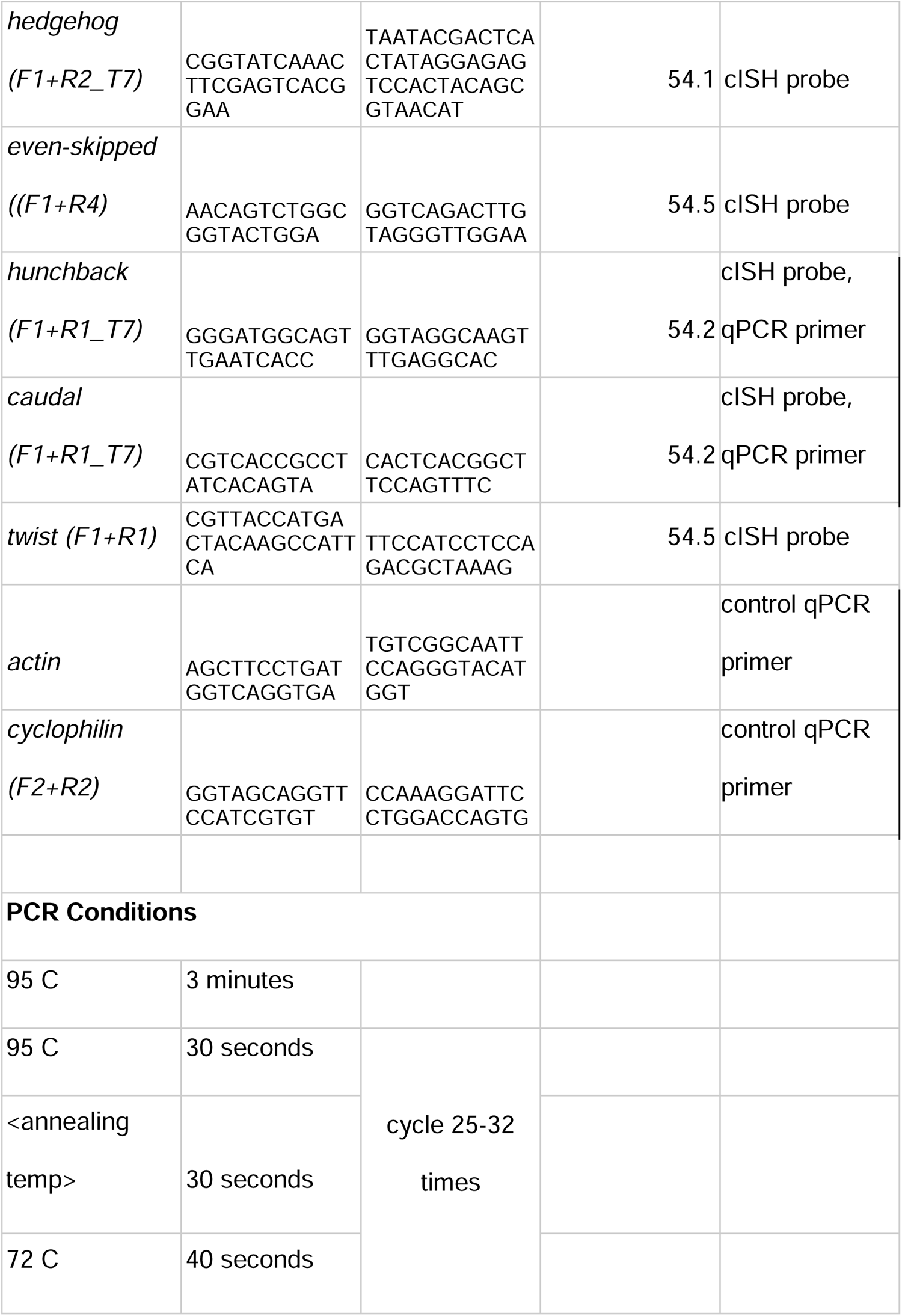
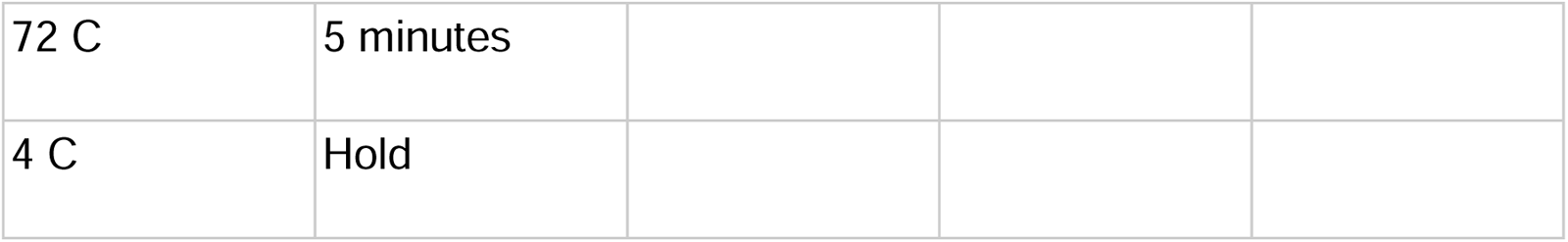
Primers and PCR conditions for gene cloning.

